# Comment on *‘Fruitless* mutant male mosquitoes gain attraction to human odor’

**DOI:** 10.1101/2021.05.24.445527

**Authors:** Brogan A. Amos, Ary A. Hoffmann, Kyran M. Staunton, Meng-Jia Lau, Thomas R. Burkot, Perran A. Ross

## Abstract

Female *Aedes aegypti* mosquitoes integrate multiple sensory cues to locate human hosts for blood meals. While male mosquitoes do not blood feed, male *Ae. aegypti* swarm around and land on humans in nature. Basrur et al. (2020) generated male *Ae. aegypti* lacking the *fruitless* gene and discovered that they gained strong attraction to humans, similar to female mosquitoes. The authors assume that host-seeking is a female-specific trait. However, all experiments were performed under confined laboratory conditions which are unable to detect long-range attraction. We used semi-field experiments to demonstrate robust attraction of male *Ae. aegypti* to humans. Our observations refute a key assumption of Basrur et al. (2020) and raise questions around conditions under which *fruitless* prevents male host-seeking. Male mosquito attraction to humans is likely to be important for mating success in wild populations and its basis should be further explored.

## Introduction

*Aedes aegypti* female mosquitoes are strongly anthropophilic (Harrington et al., 2001, McBride et al., 2014). Human skin odors, exhaled CO_2_, body heat and visual contrast all act as signals for female mosquitoes to find blood meal hosts (Liu and Vosshall, 2019). Although male mosquitoes do not blood feed, they have sophisticated olfactory systems (Wheelwright et al., 2021) used to locate female mosquitoes (Cator et al., 2009, Menda et al., 2019), nectar and other sugar sources (Barredo and DeGennaro, 2020), and conspecific males (Cabrera and Jaffe, 2007, Fawaz et al., 2014, Pitts et al., 2014).

Growing evidence demonstrates that male *Aedes* mosquitoes are attracted to humans despite it not being necessary for them to blood feed. Field observations report males swarming around and landing on humans (Banks, 1908, Hartberg, 1971, Yasuno and Tonn, 1970, Cator et al., 2011, Trpis et al., 1973, Lumsden, 1957, McClelland, 1960, Gubler and Bhattacharya, 1972). Furthermore, male *Aedes* capture rates increase when traps are baited with CO_2_ and human odor mimics (Pombi et al., 2014, Amos et al., 2020, Roiz et al., 2015, Visser et al., 2020). In a pilot experiment, Lau et al. (2020) demonstrated rapid attraction of males to humans under semi-field conditions. While males and females show similar rates of attraction to humans, sex-specific behaviors exist, with males typically swarming around humans without landing. Swarming *Ae. aegypti* males fly in a characteristic figure 8 pattern around humans. This behavior is likely to increase their reproductive success as they intercept and mate with host-seeking females (Hartberg, 1971, Cator et al., 2011, Cabrera and Jaffe, 2007).

In a recent paper, Basrur et al. (2020) claim that only female *Aedes aegypti* mosquitoes host-seek, but removal of the *fruitless* gene in males activates host-seeking behavior in male *Ae. aegypti*. Their conclusions are based on laboratory experiments, which often fail to detect male attraction to human host cues (Peach et al., 2019, van Breugel et al., 2015, McMeniman et al., 2014). The authors acknowledge some field observations but argue that they are confounded by the presence of female mosquitoes. We therefore performed experiments under semi-field conditions and demonstrate conclusively that male *Ae. aegypti* are attracted to humans in the absence of female mosquitoes.

## Results and discussion

### Male *Aedes aegypti* show long-range attraction to humans under semi-field conditions

We tested male mosquito attraction to humans under semi-field conditions using paired human-baited and unbaited traps within the same enclosure (Figure 1A). Male *Ae. aegypti* were released at a central point in the enclosure and recaptured or observed by videography at human-baited and unbaited stations. In these experiments, other objects in the semi-field cage that could potentially act as swarm markers were removed prior to initiating experiments.

**Figure 1.**
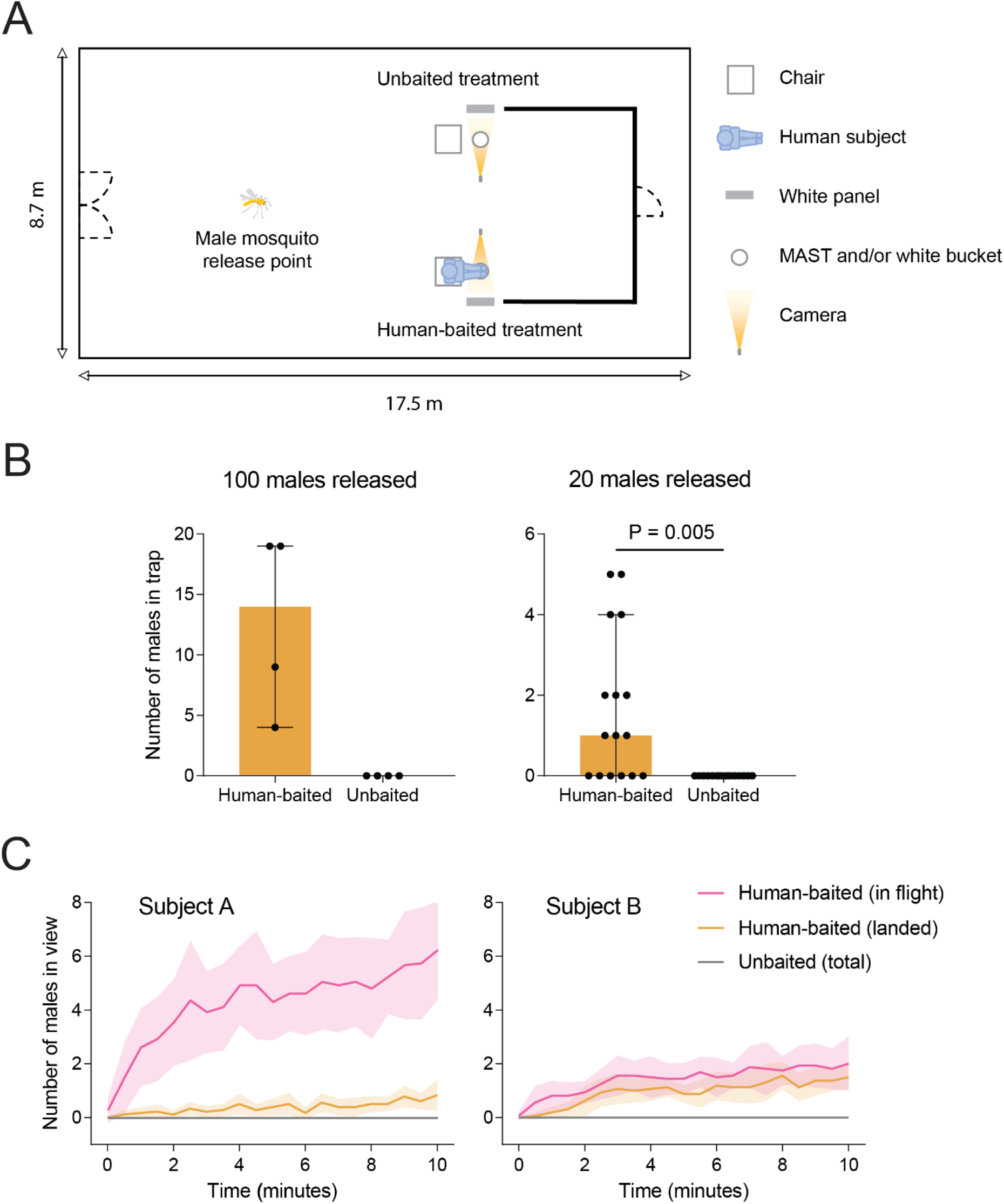
Male *Aedes aegypti* mosquitoes locate, swarm around and land on human subjects. (A) Layout of semi-field enclosure, showing the locations of human-baited and unbaited treatments for MAST (B) and videography (C) trials. (B) Number of males caught by human-baited and unbaited MASTs in 15 min when either 100 or 20 mosquitoes were released into the semi-field cage. Bars represent medians with dots showing data from individual replicate trials. Error bars are 95% confidence intervals. (C) Males in view of cameras in human-baited (colored lines) and unbaited (gray lines) treatments at 30 s intervals. Experiments were performed with two human subjects (n = 16 replicate trials per subject). Pink lines represent males in flight while orange lines show males that had landed on the human subject or the footrest. Means are shown with shaded regions representing 95% confidence intervals.

In a pilot experiment, we released 100 males (Figure 1B) and recaptured them using male *Aedes* sound traps (MASTs) (Staunton et al., 2021). After 15 min, we found that MASTs baited with a human subject sitting over the trap captured 14% (median) of males, while no mosquitoes were captured by unbaited MASTs (Figure 1B). In a second experiment using 20 males, human-baited traps captured up to 25% (median 5%) of the released males (Figure 1B). In contrast, no mosquitoes were captured by unbaited MASTs in all 16 replicates (Figure 1B). Differences in capture rates between human-baited and unbaited traps were significant according to a Wilcoxon signed-rank test (Z = 2.821, P = 0.005). Capture rates may underrepresent male attraction to humans since we observed many males swarming near the human subject that were not captured. Capture rates were higher when larger numbers of males were released, suggesting a potential effect of conspecific attraction. However, while swarming activity may increase when larger numbers of males are present, the location of males is clearly influenced by the location of the human subject.

MASTs were unable to capture many of the swarming males and the sound lures within the MASTs could plausibly influence their attraction to humans. We therefore performed a second experiment using videography to quantify male *Ae. aegypti* attraction to humans without traps. Human subjects rested their bare feet in front of a video camera facing a white plastic panel (Figure 1A). A paired treatment without a human subject was set up on the opposite side of the semi-field cage. We quantified attraction by counting the number of mosquitoes within the field of view of each camera (Video 1). Male mosquitoes began swarming almost immediately and occasionally landed on the subjects (Figure 1C), consistent with observations of male swarming in nature (Hartberg, 1971, Cator et al., 2011). The number of males observed in flight or landing increased over time, exceeding 10% after 10 min for subject A. Fewer males were viewed in flight around subject B, but differential attractiveness of humans to male mosquitoes is consistent with observations of females (Martinez et al., 2021). The number of males observed in human-baited treatments after 10 min was significantly higher than in unbaited treatments for both human subjects (Wilcoxon signed-rank test: Subject A (in flight): Z = 3.535, P < 0.001, Subject A (landed): Z = 2.751, P = 0.006, Subject B (in flight): Z = 3.077, P = 0.002, Subject B (landed): Z = 3.482, P < 0.001). Attraction rates are likely underestimated since some males within the vicinity of subjects were outside the field of view of the camera. Importantly, no mosquitoes were observed in the unbaited treatments for either subject (Figure 1C). These experiments demonstrate attraction of male *Ae. aegypti* to humans, in direct contrast to the claim by Basrur et al. (2020) that only female mosquitoes host-seek.

### Male *Aedes aegypti* mosquitoes are not attracted to humans under confined laboratory conditions

The Liverpool strain of *Ae. aegypti* used by Basrur et al. (2020) has been maintained in the laboratory for at least 80 years (Kuno, 2010) and may not be representative of wild mosquito populations. Laboratory populations of *Ae. aegypti* are typically kept in small cages with males having constant access to females, likely reducing selective pressures to maintain attraction to humans. Adaptation to laboratory conditions is therefore a plausible explanation for the lack of male host-seeking observed by Basrur et al. (2020).

We tested whether recently collected male *Ae. aegypti* from the field (F_3_ and < 6 months in the laboratory) are attracted to humans under laboratory conditions using a two-port olfactometer (30 × 30 × 30 cm, Figure 2A). Males, females or both males and females were released into a cage and collected in unbaited or human-baited traps. Females showed strong attraction to humans, with >60% being collected in human-baited traps after 5 min (Figure 2B). The number of females caught in human-baited traps was higher than in unbaited controls (Wilcoxon signed-rank test: females only: Z = 2.521, P = 0.012, females + males: Z = 2.524, P = 0.012). Rates of attraction were similar regardless of whether males were present in the same cage. In contrast to females, no males were captured in human-baited traps in any treatment (Figure 2C). Few mosquitoes were attracted to blank ports across all treatments (Figure 2B, 2C). These results are consistent with those of Basrur et al. (2020) and further demonstrate that *Ae. aegypti* show sexually dimorphic attraction to humans at close range.

**Figure 2.**
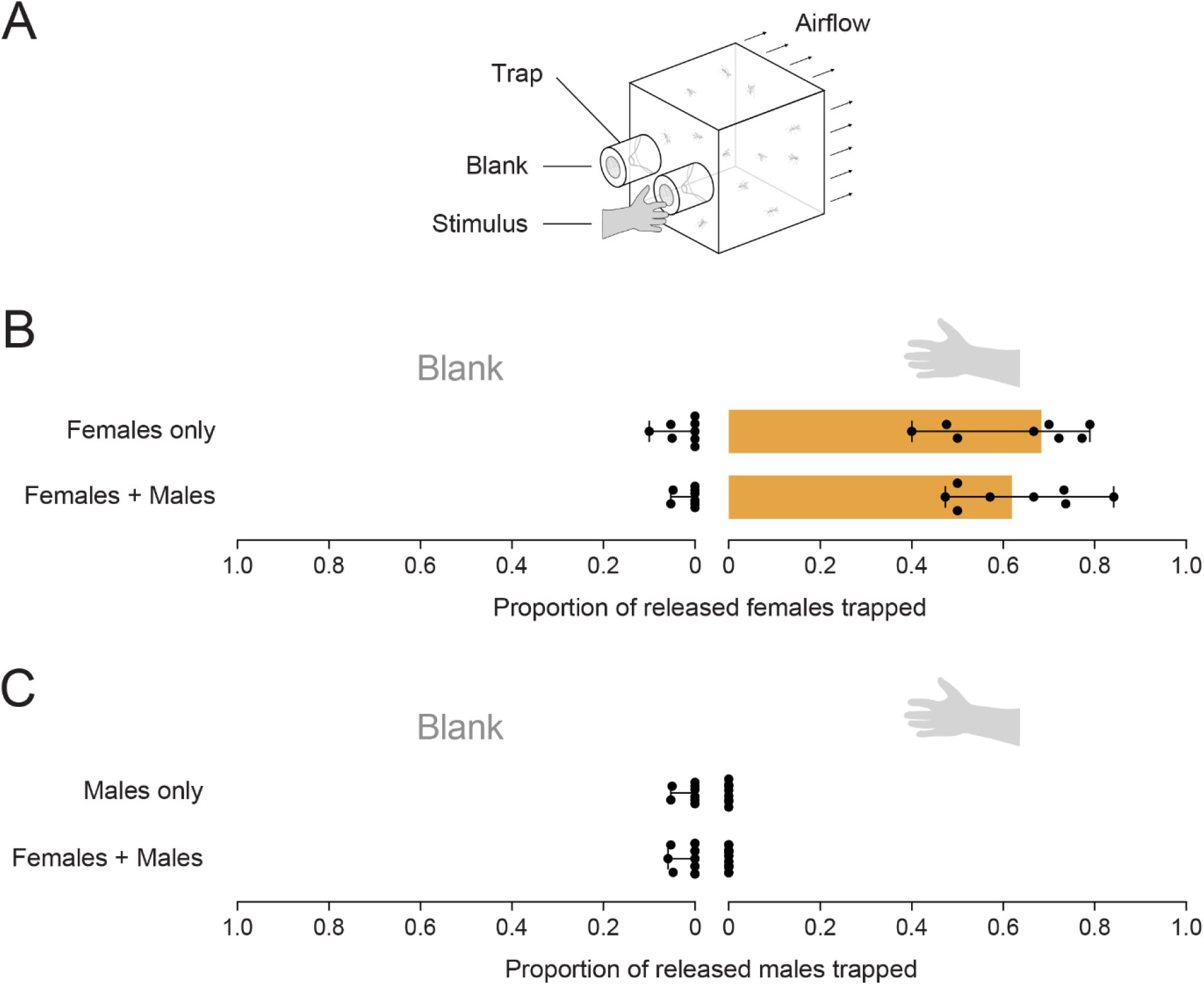
Male *Aedes aegypti* mosquitoes are not attracted to humans under confined laboratory conditions. (A) Diagram of two-port olfactometer. Mosquitoes were released into the cage and collected by one of two traps after 5 min. Traps were either unbaited (blank) or baited with the palm of a human subject (stimulus). (B-C) Proportions of released (B) females and (C) males attracted to a human hand or unbaited trap (n = 8 replicate trials each for females only, males only and females + males). Bars represent median trap proportions with dots showing proportions from individual replicate experiments. Error bars are 95% confidence intervals.

We demonstrate that male *Ae. aegypti* mosquitoes are attracted to humans in open spaces, but not under confined conditions using a port of entry assay. Our work highlights how assays performed at different scales can lead to opposing conclusions, likely because they are measuring different aspects of mosquito behavior (e.g. McMeniman et al. (2014)). In their discussion, Basrur et al. (2020) acknowledge that wild-type male mosquitoes may show some attraction to humans, but their conclusions are based on a lack of attraction in wild-type males. While we appreciate that the *fruitless* gene could contribute to sex-specific host-seeking behaviors at close range or even influence the strength of attraction to humans, our work shows that attraction to humans is already a characteristic of wild-type males. It will be interesting to explore the impact of *fruitless* in larger arenas; we predict that *fruitless* mutant males would land on humans more frequently than wild-type males under semi-field conditions.

Our study will help to inform the design of future laboratory-based behavioral assays for male *Ae. aegypti*. Further research is required to determine which host cues attract males at long distances. Male attraction to humans has important implications for mosquito control, particularly for mass-releases of males for mosquito population suppression (Crawford et al., 2020, Carvalho et al., 2015), where released male mosquitoes are likely to be regarded as a nuisance by residents in intervention areas.

Human host odors show potential as lures for traps used as vital monitoring tools in these programs, particularly in combination with sound lures.

## Materials and methods

### Mosquito strains and maintenance

*Aedes aegypti* colonies were collected from the field in Cairns (Queensland, Australia) in January 2021. Mosquitoes used in all experiments ranged from F_1_ to F_3_. Laboratory colonies were maintained at 27 ± 1°C, 70% RH with 12:12 (L:D) h regime. Adults were provided with a honey/water solution (50:50) and were blood fed using human volunteers (Human ethics approval from James Cook University H4907 and The University of Melbourne 0723847). Eggs were collected and allowed to embryonate for 3 d before being stored in air-tight containers for up to 2 mo. Eggs were hatched in water containing 0.2 g bakers’ yeast (Lowan Whole Foods, Glendenning NSW, Australia) per liter. Mosquito larvae were reared on fish food powder (TetraMin Tropical Flakes Fish Food, Tetra, Melle, Germany). Pupae were sexed and transferred to clear plastic containers (300 ml) covered with a white mesh cloth (0.5 mm pore size) with a sponge on top (30 × 40 mm^2^) soaked with honey/water solution (50:50).

### Semi-field experiments

We tested male mosquito attraction to humans under semi-field conditions though two approaches. In the first approach, we captured mosquitoes with male *Aedes* sound traps (MASTs) (Staunton et al., 2021) that were unbaited or baited with a human subject sitting near the trap entrance. In the second approach, we used videography to quantify male swarming and landing in the vicinity of a human subject, compared to an unbaited control on the other side of the cage. Experiments in semi-field cages were conducted during daylight hours in March and April, 2021. The experimental arena measures 17.5 × 8.7 m and is described in detail by Ritchie et al. (2011). Competing visual stimuli in the semi-field cage were minimized (e.g., dark-colored objects were covered with lighter-colored materials). Nitrile gloves were worn when handling objects and frequently touched objects (e.g., door handles) were regularly wiped with EtOH (80%) throughout the experimental period to minimize human odor interference. Male mosquitoes used in semi-field experiments were unmated, between 2- and 7-d post-emergence. All human subjects acting as lures wore light-colored clothing, minimized movement, and refrained from using perfumed products 24 h before and during the trials.

#### MAST trials

MASTs use sound frequencies which mimic female mosquito flight tones to capture male *Ae. aegypti* (Staunton et al., 2021). MAST trials involved two paired treatments within the same semi-field cage (Flight cage B). MASTs were cleaned with EtOH (80% aqueous solution) to remove potential human skin odors before use and thereafter only handled with nitrile gloves. The black MAST bases (which act as swarm markers) were not used in these trials. Instead, MAST heads (the capture container components of the trap system which include the sound lures) were placed on upturned white plastic buckets (3L) such that they were 15 cm above the ground. MASTs were placed 5.5 m apart outside of a structure built to resemble the downstairs area of a traditional house in Queensland. A white plastic and metal chair was placed over each trap. A human (subject A in the below experiment) acting as bait sat in one chair, with the other left empty, with the positions of human-baited and unbaited treatments swapping each replicate. Male *Ae. aegypti* were released remotely at a central location in the cage approximately 6 m from the traps. We ran 4 replicate trials with 100 males at sound lure settings of 495 Hz, continuous tone (volume level 1), and 16 replicate trials with 20 males at sound lure settings of 550 Hz, continuous tone (volume level 2). Trials ran for 15 min and mosquitoes captured by the MAST were counted, with an equal number of new males released between each replicate. Numbers of males captured by human-baited and unbaited MASTs were compared using Wilcoxon signed-rank tests. All data were analyzed using SPSS statistics version 24.0 for Windows (SPSS Inc, Chicago, IL).

#### Videography

Videography trials involved two paired treatments on opposite sides of flight cage A (3.2 m apart) (Figure 1A). Two cameras (Professional Series Motorised Bullet 8MP cameras (VIP Vision™)) were installed at ground level facing upturned plastic white (3L) buckets in front of white corrugated plastic panel (600 × 800 × 5 mm; Corex Plastics Australia Pty. Ltd.). In the human-baited treatment, a human subject sat on a white plastic and metal chair and placed their bare feet on the bucket for the duration of each trial. Unbaited treatments were set up identically to human-baited treatments, but on the opposite side of the cage and without a human subject. Two subjects were used, subject A (female, Caucasian, age 32) and subject B (male, Caucasian, age 29), and the position of human-baited and unbaited treatments was swapped each replicate (n = 16 replicate trials per subject). Before trials commenced, 50 male *Ae. aegypti* were released remotely at a central location in the cage once per day. Video footage was used to count the number of visible mosquitoes (both flying and landed in the frame (approximately 1200 × 700 mm field of view, covering the vicinity of the subject’s feet and lower legs in the human-baited treatment) every 30 sec for 10 min, starting at time zero when the participant placed their feet on the bucket. Numbers of males observed in human-baited and unbaited treatments at 10 min were compared using Wilcoxon signed-rank tests.

### Laboratory olfactometer assays

We compared the attraction of male and female *Ae. aegypti* to a live human host (male, Caucasian, age 31) under confined laboratory conditions. Experiments were performed in a two-port olfactometer (30 × 30 × 30 cm, Figure 2A) identical to the one used by Ross et al. (2019), except that the stimulus ports were removed. We performed three treatments: males only, females only and females + males, with each treatment replicated eight times. Mosquitoes of both sexes were 6-7d post-emergence, mated, and sugar-starved for approximately 24 hr. In each treatment, approximately 20 adults per sex were released into the cage and left to acclimate for 1 min. A box fan placed at the opposite end of the cage drew air (∼0.2 m/s) through two traps into the cage. The hand of a human subject was placed 1 cm in front of one of the traps, with the other blank. Sides were alternated each replicate. After 5 min, the entrances to both traps were closed and the number of males and/or females in each trap as well as the cage were counted. Mosquitoes that were damaged before or during the experiment were excluded.

Proportions of males collected in stimulus and blank traps after 5 min were compared using Wilcoxon signed-rank tests.

## Acknowledgements

We thank Tom Swan for assistance with the videography experiment and Verily for providing consent to use the male *Aedes* sound traps in this study. We also thank Nipun Basrur, Leslie Vosshall and Conor McMeniman for providing valuable feedback on the first version of this manuscript.

## Video legends

**Video 1. Video evidence of male *Aedes aegypti* attraction to humans**. Male *Ae. aegypti* are shown swarming around the legs of human subject A (left) in representative footage from the videography experiment. No activity is seen in the unbaited control (right). Videos have been cropped for clarity; unedited footage from treatments and controls of both subjects are provided in Videos S1-S4.

## Supplementary information

**Video S1**. Representative footage from the human-baited treatment for subject A in the videography experiment. This footage was used to quantify male *Aedes aegypti* attraction to humans in Figure 1C.

**Video S2**. Representative footage from the unbaited treatment for subject A in the videography experiment. This footage was used to quantify male *Aedes aegypti* attraction to humans in Figure 1C.

**Video S3**. Representative footage from the human-baited treatment for subject B in the videography experiment. This footage was used to quantify male *Aedes aegypti* attraction to humans in Figure 1C.

**Video S4**. Representative footage from the unbaited treatment for subject B in the videography experiment. This footage was used to quantify male *Aedes aegypti* attraction to humans in Figure 1C.

